# *N*-glycosylation of CD24 mediates cell motility but inhibits cell proliferation in colorectal cancer

**DOI:** 10.1101/2021.05.02.442315

**Authors:** Zaki Hakami, Teresa Raposo, Abdullah Alsulaiman, William Dalleywater, Hassan Otifi, Charlotte Horobin, Karin Garrie, Mohammad Ilyas

**Author notes:** Correspondence to Prof. Mohammad Ilyas: West block A floor, Pathology Department, Queen’s Medical Centre, Derby Road, NG7 2UH Nottingham, United Kingdom.

## Abstract

CD24 confers features of stemness and enhances cell motility in colorectal cancer. It contains numerous *O*-glycosylation sites but just two putative *N*-glycosylation sites. The study aimed to mutate *N*-glycosylation as it thought to be important in mediating many glycoproteins’ function.

Site-directed mutagenesis performed to mutate *N*-glycosylation sites by converting asparagine residues to glutamine. Each site was mutated individually (CD24^N36Q^ and CD24^N52Q^) and in combination (CD24^N[36,52]Q^) and were overexpressed in HCT116 and SW480 cell lines. The effect of the mutant CD24 on cell viability, cell motility and induction of downstream targets was compared to wild-type CD24 and empty vector control (EVC).

Overexpression of the individually mutated clones increased cell motility compared to EVC but was significantly less than that induced by CD24^WT^. Overexpression of the double mutant clone resulted in near-complete abrogation of cell motility induction. Unexpectedly, the expression of mutant clones (single and double) increased cell viability. Analysis of downstream targets and F-actin staining demonstrated reduced induction of known CD24 targets by mutant vectors and reduced epithelial-mesenchymal transition.

In conclusion, each *N*-glycosylation site on CD24 contributes partially to induce motility by CD24, and these are additive. Loss of the *N*-glycosylation results in an unexpected gain of proliferative function.

**Highlights:**

- CD24 is a heavily glycosylated molecule that contains two N-glycosylation sites, and these contribute to the activation of downstream targets and induction of cell motility
- Each N-glycosylation site contributes partially, and their effects appear to be additive
- The N-glycosylation sites may exert an inhibitory effect on cell proliferation
- Much of the CD24 activity is mediated through post-transcriptional effects

## Introduction

Colorectal cancer (CRC) is the third most common cancer globally, and it is the second most common cause of mortality [1]. Advancements in diagnosis and treatments have shown an improvement in overall survival (OS) rates for early CRC stages; however, 20% of cases present with metastasis and a further 35-45% have metastatic recurrence [2], leading to a 5-year OS of <10% [3].

Tumour metastasis follows distinct steps through epithelial-mesenchymal transition (EMT), a process in which epithelial cells undergo biological changes, including a loss in cell polarity, disruption of cell-cell adhesion and restructuring of the actin cytoskeleton. These changes contribute to the acquisition of invasive and migratory tendencies [4]. We have previously shown that CD24 induces cell invasion and migration in CRC, and it is therefore, a good candidate for a “metastasis-inducing” gene [5]. The invasion and migration activities are mediated through a CD24–ILK/FAK signalling pathway (unpublished data) and are accompanied by a “cadherin switch”, where expression of E-cadherin is reduced and N-cadherin expression, characteristic of mesenchymal cells, is increased [6]. In multiple types of tumour (such as breast cancer [7], gastric cancer [8], non-small cell lung cancer [8], ovarian cancer [9] and pancreatic cancer [10]), CD24 is associated with promoting cancer cell migration and invasion and with reduced OS [11,12].

CD24 gene is located on chromosome 6q21, and it is a highly glycosylated, glycosyl-phosphatidyl-inositol-anchored (GPI) protein [13]. Human mature CD24 comprises 31 amino acids, nearly half of which are serine and threonine residues rendering it a mucin-like protein [14]. Analysis of the amino acid sequence shows that CD24 has 16 potential *O*-glycosylation sites and two potential *N*-glycosylation sites [13]. Glycosylation is a post-translational modification involving the enzymatic glycosidic linkage reaction between the anomeric hydroxyl group of sugar and a protein [15]. During the glycosylation process, carbohydrates can be attached to proteins and lipids. For *N*-glycosylation, carbohydrates are attached to an asparagine residue that appears in a specific sequon. Studies have shown that *N*-linked glycosylation’s target sequence is asparagine-(X)-serine/threonine, where X can be any amino acid except for proline [16].

In normal growth and development, *O*-glycosylated residues have structural roles. In contrast, *N*-glycosylated residues have functional roles and have been shown to play a role in protein biology, including folding, secretion, stabilisation, conformation and trafficking to cellular and subcellular sites [17]. In cancer, *N*-glycosylated proteins are reported to play a role in tumour growth and development by way of enhancing the cancer cell’s ability to evade immune detection, promoting uncontrolled proliferation, migration, and cell-cell adhesion [18-20].

For this study, we hypothesised that the *N*-glycosylation sites promote CD24-induced motility by regulating EMT and thereby, they may have a role in cancer metastasis. To test this, site-directed mutagenesis (SDM) was performed to mutate the two putative *N*-glycosylation sites in CD24 by substituting asparagine with glutamine. Mutant clones, including CD24 wild-type and EVC, were overexpressed in two CD24 negative CRC cell lines, HCT116 and SW480 [21]. Following cell line transfection, functional studies *in vitro* and protein/RNA analysis were performed.

Our preliminary data in this study confirmed that the N36 and N52 sites are bona-fide N-glycosylation sites. In addition, this study indicated that the ability of CD24 to regulate cell motility, EMT and cadherin switch in CRC cells was dependent on N-glycosylation. These data suggest that targeting the N-glycosylation sites present in CD24 may help improve prognosis and may be used as a targeted therapy against CRC.

## Materials and Methods

### Cell Culture

HCT116 and SW480 cell lines were obtained from the American Type Culture Collection (ATCC). Cells were cultured in Dulbecco’s modified Eagle’s medium (DMEM, GIBCO®, Invitrogen) supplemented with 10% fetal bovine serum (FBS, Sigma) and 1% L-glutamine (Sigma), incubated in a humidified 5% CO_2_ environment at 37°C. The cell lines were regularly passaged every 3-4 days by dissociation with Trypsin-EDTA 0.25% (Sigma Aldrich). The cell lines were regularly subjected to monthly screening for mycoplasma infection. The identity of the cell lines was also confirmed by short tandem repeat (STR) genotyping using PCR-single locus technology.

### Site-Directed Mutagenesis (SDM)

The Asparagine to Glutamine (N-Q) point mutation was performed using a Phusion SDM kit (Thermo Fisher Scientific, CAT# F541). In this assay, the mutation was introduced at each of the *N*-glycosylation sites (i.e., N36Q and N52Q) individually in a CD24 expression construct. Briefly, the PCR mix consisted of 10µL Phusion HF buffer, 200µM dNTPs, 0.5µM each of forward and reverse primer **(Table S1)**, 1ng/µL template DNA, 0.02U/µL Phusion hot star II DNA polymerase (2U/µL), made up to 20µL with nuclease-free water. Using the Primus 96 Thermal Cycler (MWG-Biotech Inc., Ebensburg, Germany), the cycling protocol was as follows: X1 initial denaturation cycle at 98°C for 30-sec, X25 cycles of (denaturation at 98°C for 10 sec, annealing at 67-68°C for 30 sec per primer, extension at 72°C for 3min), X1 final extension cycle at 72°C for 10min. The products were run on 1% agarose gel, and the fragments purified using a GenElute™ Gel extraction Kit (Sigma-Aldrich). The PCR products were circularised with T4 DNA ligase following the manufacturer’s instructions (Thermo Fisher Scientific). 10-20ng (made up to 5 μl) of DNA plasmid was added to 10µL ligation reaction mixture (2.5 μl nuclease-free water, 2 μl rapid ligation buffer, and 0.5 μl T4 DNA ligase enzyme) and incubated at room temperature for 5min. Ligation control mixture was prepared without T4 DNA ligase. According to standard protocol, the ligated products were transformed into NEB 5α competent cells (New England BioLabs) using the heat shock method. The plasmid was extracted using a mini-prep extraction kit (Sigma-Aldrich), and the introduced mutations were confirmed by DNA sequencing.

To generate a double mutant plasmid, the same protocol was used as mentioned above. However, the constructed single mutant plasmids, either N36Q or N52Q, were used for SDM instead to generate a construct containing a mutation at both sites (CD24^N[36,52]Q^).

### Transient Transfection

The transfection protocol was as follows; 5×10^5^ cells were seeded into 6-well plates containing 2mL fresh growth media and incubated for 24h. The medium was discarded, and the cells were serum-starved for 1h by adding 1.5mL Opti-MEM® reduced-serum medium (Gibco, Life Technologies) per well prior to transfection. During incubation, the transfection mixture was prepared as follows: 8µL/well of Lipofectamine™ 2000 was added to 250µL Opti-MEM® and incubated at room temperature for 5min. In a separate tube, 4µg/well of CD24 mutant plasmids or empty vector control were added to 250µL Opti-MEM®, and the two tubes were then combined and incubated for 20min at room temperature. The plasmid and Lipofectamine mixture was added to the well plates, incubated for 4-6 h. After incubation, the transfection media was replaced with 2 ml fresh growth media and the cells were harvested 24-48h post-transfection for protein extraction and *in vitro* functional assays. The transfection efficiency was determined according to the protocol provided by the Flow Cytometry Facility at the University of Nottingham **(Method S1)**.

### Protein extraction and Western Blotting

Whole-cell lysates were used for Western blotting. Protein extraction occurred following cell lysis by adding 150 µL Pierce® radioimmunoprecipitation assay (RIPA) buffer (Thermo Fisher Scientific) and 1.5 µL protease and phosphatase inhibitor cocktail (10X Thermo Fisher Scientific) in a dilution of 1:100. Protein quantification was performed using a Pierce™ BCA kit (Thermo Fisher Scientific). For Western blotting, 50µg of protein samples were added to loading buffer (one-third volume of protein sample) containing 1x NuPAGE™ sample reducing agent (10x, Thermo Fisher Scientific) denatured at 95°C. Protein fractionation occurred by running samples on a NuPAGE 4-12% Bis-Tris gel (Invitrogen) at 150V for 2h. Protein was transferred to a polyvinylidene difluoride (PVDF) membrane (Amersham, GE Healthcare Life Sciences) by using a semi-dry transfer on Bio-Rad Transblot turbo transfer system (Bio-Rad, Watford, UK). The membrane was blocked for 1hr with 5% skimmed milk (Sigma-Aldrich) in Tris-buffered saline with 0.1% Tween 20 (TBST) at room temperature, then incubated overnight at 4°C with primary antibodies: SWA11 (donated by Professor Altevogt, Germany), E-cadherin [#3195), N-cadherin [#13116], Snail [#3879], vimentin [#5741] (all from Cell Signaling Technology) and α-tubulin (Abcam, Ab7291) **(Table S2)**. The membrane was washed with TBST then incubated with appropriate horseradish peroxidase-conjugated secondary antibody (Sigma-Aldrich, Anti-mouse CAT# A4416 and Anti-Rabbit #A6154) for 1h at room temperature. The membrane was rewashed with TBST, and the visualisation occurred using enhanced chemiluminescence (ECL) detection kit (Thermo Fisher Scientific) on the Odyssey Fc imaging system (LI-COR Bioscience, UK). Protein expression was digitally calculated using the Image Studio-Lite Ver 5.2 software (LI-COR Biosciences Ltd. UK). Protein expression was normalised by calculation of the ratio between target proteins to the housekeeper α-tubulin. The N-glycosylation digestion was performed according to the manufacturer’s protocol **(Method S2)**.

### RNA Extraction and Quantitative PCR

Total RNA was extracted using an RNA extraction kit (Sigma-Aldrich) according to the provided manual. 1μg to 2μg of RNA was reversed transcribed to cDNA using M-MLV reverse transcriptase following the manufacturer’s protocol **(Method S3)**. RT-qPCR was performed to determine the mRNA levels of CD24 and EMT biomarkers **(Table S3)** using GoTaq® qPCR Master Mix with SYBR green (Promega). The qPCR was performed in a total volume of 20µL per-reaction containing: 10ng of cDNA, 0.250µM each of the forward and reverse primers, 10µL of GoTaq® qPCR Master Mix and made up to 20µL with nuclease-free water. The PCR was run on the following thermal cycles: X1 cycle hot-start activation for 2 mins at 95°C, X40 cycles (denaturation at 95°C for 15 sec, annealing at 60°C for 60 sec, extension at 72°C for 60 sec), X1 cycle melting curve stage (95°C for 60 sec, 55°C for 30 sec, 95°C for 30 sec). Relative mRNA levels of target genes in experimental conditions were normalised to porphobilinogen deaminase (PBGD, housekeeping control) and the empty vector control using Livak’s comparative threshold cycle (2-ΔΔCT) method [22].

### Cell Proliferation

A cell viability assay was used to quantify cell numbers. Five thousand cells were seeded into each well of 96-well plates containing 100µL growth medium and incubated 24h. The media was removed and washed twice with phosphate-buffered saline (PBS). The PrestoBlue® reagent (Invitrogen) was diluted in 10% FBS–DMEM (dilution of 1:10), and 100 µl of the mixture added to each well for 30min. Fluorescence quantification of resazurin product (resorufin) was performed using the BMG FLUOstar OPTIMA plate reader (Ortenberg, Germany; FlexStation II, Molecular Devices, San Jose, CA, USA) at 544nm excitation and 612nm emission wavelength. Post-reading, the PrestoBlue® mixture was discarded, cells were washed once with medium and then replaced with fresh growth medium, and incubated for a further 24h. Cell proliferation was evaluated at 0, 24, 48, and 72h. Relative fluorescence unit (RFU) readings were normalised to the reading obtained at 0hr and displayed relative to cell viability.

### Invasion and Migration Assays

Cell migration was measured using both the transwell migration assay and the wound healing assay, whilst cell invasion was measured by transwell motility through Matrigel. Assays were performed in triplicate on at least two separate occasions. Transwell migration was measured using 8.0-μm pore size Transwell polycarbonate membrane inserts (Costar, Corning). Firstly, 650µL of fresh medium (containing 10% FBS) was added to the lower receiver wells as a chemoattractant. To the upper inserts, 1×10^5^ cells were seeded in 100 µl DMEM (containing 0% FBS) and incubated at 37°C in 5% CO2. Cell migration was assessed after 24h by incubation of the inserts for 5min in a new well-containing 650µL trypsin-EDTA to detach the cells. Cells were then transferred to the original well and labelled with 3μM calcein acetoxymethyl ester (Thermo Fisher Scientific) by incubation for 1hr at 37°C. Cell invasion was measured in the same way except that before seeding, the insert was prepared by coating in 300μg/ml Matrigel (Matrigel® Matrix, Corning®) diluted in DMEM (0% FBS) and allowed to polymerise for 2h at 37°C in 5% CO_2_. Visualisation of the cell motility was performed using Nikon Eclipse TiE fluorescence microscope (fluorescein isothiocyanate filter, 488nm excitation). Four random microscopic fields per well were taken at ×10 magnification, and the cells counted manually using ImageJ (National Institutes of Health, Bethesda, MD) and analysed by taking the average of the four fields per well **(Figure S3)**.

### Wound Healing Assays

Wound healing was assessed using Culture-Insert 2-Well 24, ibiTreat (ibidi) placed in the centre of each well in 24-well plates. Following the manufacture’s protocol, 1×106 cells were suspended in 1ml DMEM (0% FBS), 70µL/well was added to the insert and incubated for 24h. Following this, the inserts were removed, the cell layers were washed with PBS to remove non-adherent cells and covered by 1ml fresh medium. Wound closure was visualised at 0h and 24h using a wide-field system Okolab microscope. Cell migration across the gap was measured by calculating the surface area of the wound using ImageJ. In this assay, 10 μg/ml mitomycin C (Sigma-Aldrich) was added to inhibit cell proliferation.

### F-actin staining

Filamentous actin (F-Actin) was visualised by immunofluorescence. Cells were seeded at 2×10^4^cells/well in 8 well chamber slides (Corning) and incubated for 24h. They were fixed with a mixture of 4% paraformaldehyde and 5% sucrose for 10mins and permeabilised using 0.1% Triton X-100 in PBS (Sigma-Aldrich) for 5min at room temperature. Cells were stained by incubating for 1 hour with Texas Red-X phalloidin (Thermo Fisher Scientific) in 100 μl of 3% bovine serum albumin (BSA, Fisher Scientific) diluted in PBS. The nuclei were stained by incubation with 4′,6-diamidino-2-phenylindole (DAPI; Sigma-Aldrich) diluted 1:10,000 in 1% BSA for 15min, and slides were mounted with Fluoroshield anti-fade mountant (Sigma-Aldrich). The slides were imaged using a confocal Leica DMI4000B microscope at x20 magnification.

### Statistical Analysis

Statistical analysis was performed using GraphPad Prism version 7.03 (La Jolla, CA). All data were evaluated using a two-way analysis of variance (ANOVA) to analyse cell proliferation, migration, and invasion. The mean protein expression in the western blotting was compared using the Student’s *t*-test or one-way ANOVA. The value of p<0.05 was considered statistically significant.

## Results

### Mutant constructs show overexpression of CD24

The successful creation of the CD24^N36Q^, CD24^N52Q^, and CD24^N[36,52]Q^ expression constructs was confirmed by direct sequencing **(Figure S2)**. The mutant expression constructs, alongside expression constructs for wild-type CD24 (CD24^WT^) and empty vector controls (EVC), were transfected into HCT116 and SW480 cells. Western blotting also confirmed that mutant protein from all the constructs was successfully expressed in both cell lines **(Figure 1 A, B)**. It is well known that heavily glycosylated proteins such as CD24 produce smeared bands on Western blotting, and the appearance may be influenced by both the pattern and amount of glycosylation. Since we altered putative N-glycosylation sites, a change in the banding pattern would be expected in the mutant protein if they were true glycosylation sites. However, assessment of expression levels and protein size is not reliable in Western blots of glycosylated proteins. To ensure that the banding alterations were not due to changes in expression of the core protein caused by unidentified mutations in the plasmid, the lysates were digested using an N-glycosylation digestion mix (α2,3,6,8 Neuraminidase and PNGase F) to remove all N-linked oligosaccharides prior to Western blotting. A single 10kDa band was visualised in all conditions, and similar levels of both mutant and wild type proteins were shown to be expressed **(Figure 1C, D)**. This confirmed that any changes seen were due to changes in glycosylation rather than changes in protein expression.

**Figure 1.**
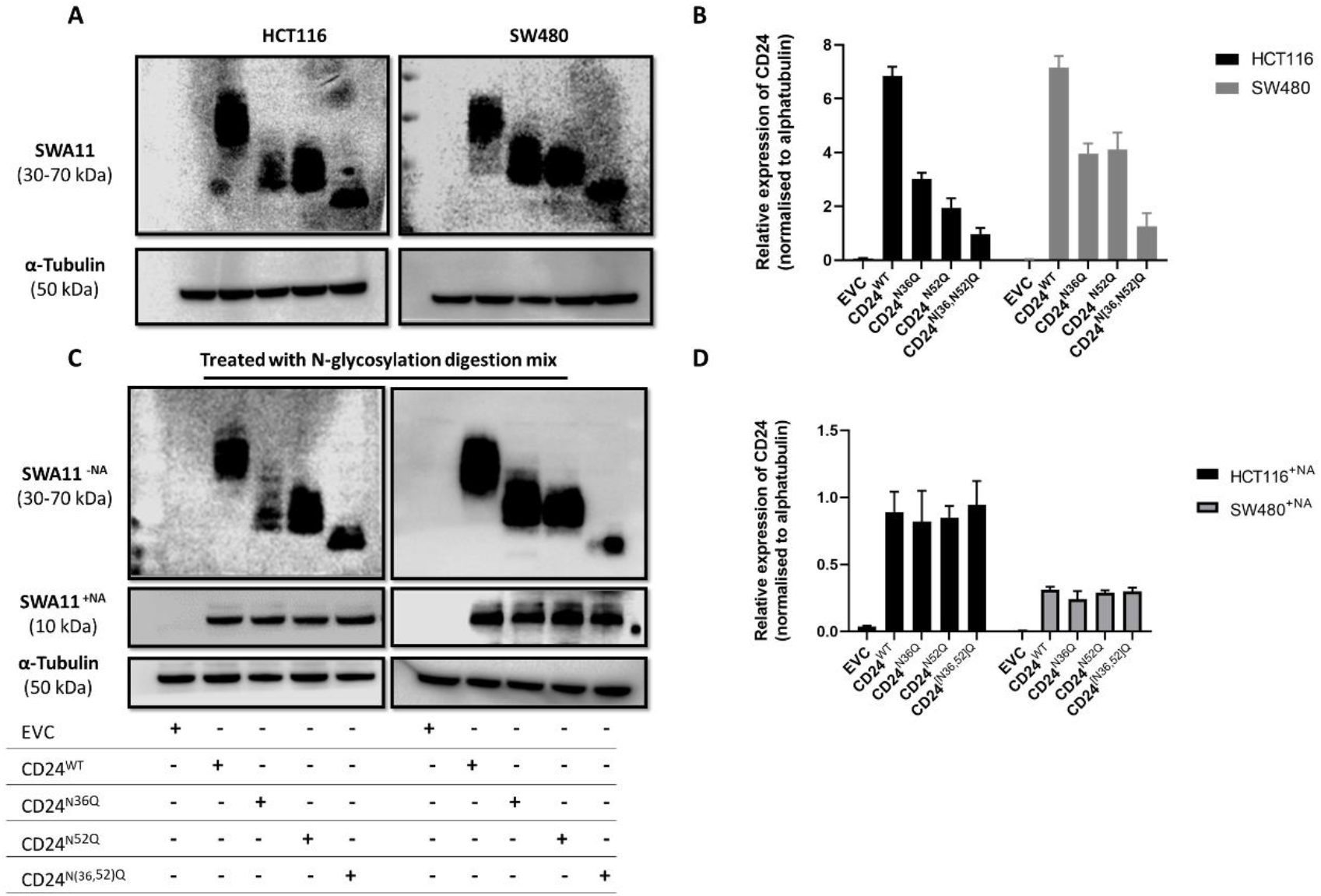
The expression level of CD24 shows variable glycosylation in mutant constructs. All mutants constructs blotted with SWA11 (CD24) antibody were successfully overexpressed in both HCT116 and SW480 with bands variation compared to CD24^WT^ in Western blot. The cropped blots are used in the figure. The samples derive from the same experiment and that the blots were processed in parallel (A). Densitometry and quantification of western blot normalised to alpha-tubulin (B). After N-glycosylation digestion, the relative protein expression of CD24 in HCT116 and SW480 cells expressing CD24^WT^ and CD24 mutant constructs showed similar bands at approximately 10kDa (C). The bar graph (D) represents the quantitative densitometry analysis of western blot for CD24 protein normalised to alpha-tubulin. The figure displayed a combined result of the three experimental replicates. EVC was used as negative control and CD24^WT^ as a positive control. Abbreviation: EVC; empty vector control, -NA, with (Neuraminidase+PNGase F), +NA, without (Neuraminidase+PNGase F).

### Loss of CD24 *N*-glycosylation sites results in increased cell proliferation

Transfection efficiency can be assessed by both Western blotting and flow cytometry. Evaluation by flow cytometry showed that all mutant constructs and CD24^WT^ had the same efficiency of approximately 50% to 70% **(Figure 2A, B)**. Our previous studies on the effect of CD24 on cell proliferation have shown that CD24 had no role in cell proliferation [23]. In order to investigate whether the *N*-glycosylation sites of CD24 play a role in regulating cell proliferation, a PrestoBlue cell proliferation assay was used to determine proliferation at four consecutive time points **(Figure 2 C, D)**. As expected, there was no difference in cell numbers between cells transfected with CD24^WT^ and cells transfected with EVC (i.e., CD24^WT^ vs EVC) in either HCT116 or SW480. Unexpectedly, transfection with the single-site mutant constructs CD24^N36Q^ and CD24^N52Q^ (i.e., CD24^N36Q^ vs CD24^WT^ and CD24^N52Q^ vs CD24^WT^) resulted in a significant increase in cell numbers from 24 hours onwards in both cell lines (p<0.0493 at 24h and p<0.001 at the other time points). The size of the effect was similar in each mutation, and no statistical difference was observed for CD24^N36Q^ vs CD24^N52Q^. However, when the cells were transfected with the double mutant construct, the cell number was significantly greater than that seen in single mutant transfections, i.e. CD24^N[36,52]Q^ vs CD24^N36Q^ and CD24^N[36,52]Q^ vs CD24^N52Q^ (p <0.0001 at each time point). The observations were similar in both cell lines, and the greatest effect was seen in the comparison of the double mutant with wild type, i.e. CD24^N[36,52]Q^ vs CD24^WT^, p <0.0001 at each time point for both HCT116 and SW480.

**Figure 2.**
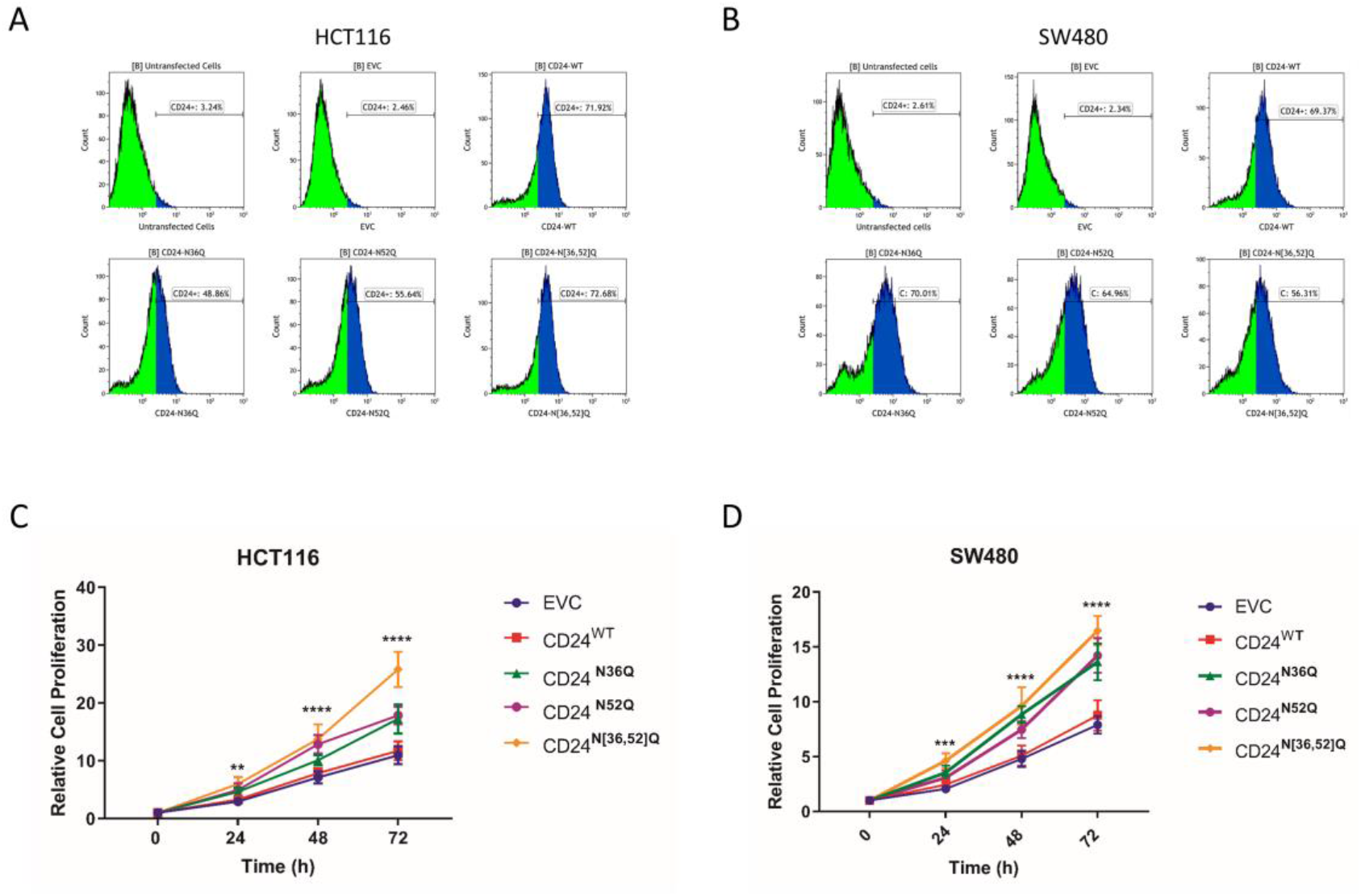
Mutations in the N-glycosylation sites of CD24 influences cell proliferation. The percentage of transfection efficiency for HCT116 (A) and SW480 (B) stained with CD24+ was indicated for each construct, and the non-transfected cells and EVC were used as negative controls. The cells population that shifts to the right of the curve are CD24+ cells. Relative cell proliferation values of HCT116 (C) and SW480 (D) cell lines show a significant increase in cell viability for all mutants compared to the EVC and CD24WT at 24hrs (***P≤0.001), 48hrs and 72hrs (****P≤ 0.0001, respectively). Results are representative of three biological replicates, and error bars indicate the SD.

### Loss of CD24 *N*-glycosylation sites results in loss of motility induction

We have previously shown that CD24 is able to induce cell motility. Examinations into the potential role of *N*-glycosylation in cell migration were performed by comparative transwell migration assays and wound healing assays using cells transfected with mutant constructs and CD24^WT^. As expected, and in concordance with our previous data, CD24^WT^ vs EVC demonstrated a marked increase in transwell cell migration in HCT116 and SW480 (P<0.0001 for each), **(Figure 3A)**. Transfection of the single mutant plasmids (CD24^N36Q^ vs EVC and CD24^N52Q^ vs EVC) showed increased cell migration, but this was significantly less than that induced by CD24^WT^ (CD24^N36Q^ vs CD24^WT^ and CD24^N52Q^ vs CD24^WT^, P= 0.0018 for HCT116, p= 0.0045 for SW480). Transfection with the double mutant CD24^N[36,52]Q^ caused further reduction in cell transwell migration compared to CD24^WT^ (CD24^N[36,52]Q^ vs CD24^WT^, P<0.0001 in both HCT116 and SW480) **(Figure 3A)**. The double mutant retained some ability to induce motility compared to the EVC in HCT116 (CD24^N[36,52]Q^ vs EVC, P< 0.0114), but this appeared to be lost in SW480 (Figure 3, CD24^N[36,52]Q^ vs EVC, p= NS).

**Figure 3.**
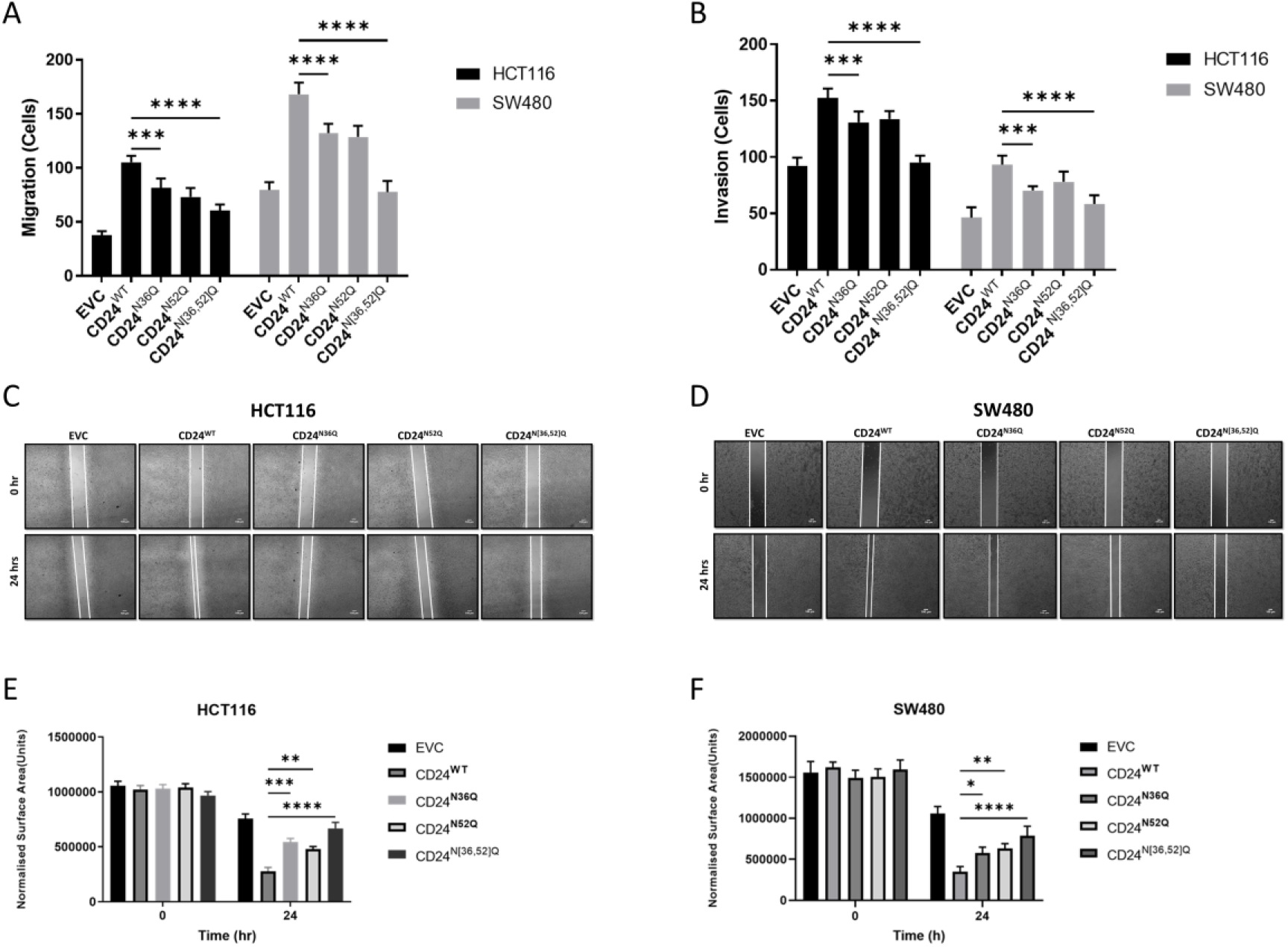
N-glycosylation mutation causes inhibition in CD24-induced cell invasion and migration. Transwell migration and invasion assays of HCT116 and SW460 cells transfected with CD24^N36Q^ and CD24^N52Q^ shows a partial reduction compare to the control CD24 whilst assays CD24 ^N[36,52]Q^ cells show significant depletion in both assays **(A, B)**. Wound healing assay of HCT116 cells **(C)** and SW480 **(D)** showed that the overexpression of CD24WT reduces the wound gap compared to EVC. This ability was inhibited when both CD24 N-glycosylation sites are mutated. The lower panel graphs represent the normalised wound gap area of HCT116 (Left) and SW480 (Right). * P < 0.05, ** P < 0.01, *** P < 0.001, ****P>0.0001.

Cell motility was also tested using a wound-healing assay in which residual wound size was measured over 24 hours (as described in material and methods). The data in **(Figure 3C, D)** replicated those seen in the transwell migration assay inasmuch as the residual wound was significantly smaller in size in CD24^WT^ vs EVC in both cell lines (HCT116 p<0.0001, SW480 p<0.0001). The residual wound size was significantly smaller in CD24^N36Q^ vs EVC (HCT116 p> 0.0046, SW480 p<0.0072) and CD24^N52Q^ vs EVC (HCT116 p<0.0002, SW480 p<0.0045) but the residual wound was significantly smaller in CD24^WT^ vs CD24^N36Q^ (HCT116 p<0.0242, SW480 p<0.0181 and CD24^WT^ vs CD24^N52Q^ (HCT116 p<0.0087, SW480 p<0.0367). Consistent with the transwell migration data, the difference in residual wound size was even more marked for CD24^N[36,52]Q^ vs CD24^WT^ (P<0.0001 for HCT116 and SW480, respectively) and wound healing was almost the same in CD24^N[36,52]Q^ vs EVC for both HCT116 and SW480.

Our previous data showed that CD24 could also induce cell invasion through Matrigel (unpublished for cell invasion). Comparative transwell invasion assays performed examinations into the potential role of N-glycosylation in cell invasion. In concordance with our previous data, CD24^WT^ vs EVC demonstrated a marked increase in invasion in HCT116 and SW480 (P<0.0001 for each) **(Figure 3B)**. As with the migration assays, single mutant plasmids showed increased cell invasion than EVC (CD24^N36Q^ vs EVC and CD24^N52Q^ vs EVC) in both cell lines but were less significant than CD24^WT^ (P=0.0012). Double mutants CD24^N[36,52]Q^ caused further reduction and near-total abrogation in invasion (CD24^N[36,52]Q^ vs CD24^WT^, P<0.0001 in both HCT116 and SW480. No significant difference was observed in CD24^N[36,52]Q^ vs EVC for both cell lines.

### Loss of CD24 *N*-glycosylation sites results in loss of activation of downstream targets and reduction in epithelial-mesenchymal transition (EMT)

Previously in our lab, CD24 was shown to promote EMT [6]. To investigate the potential role of N-glycosylation on EMT, HCT116 and SW480 were transfected with CD24^WT^ and CD24^N[36,52]Q^, and the expression of EMT-related markers (E-cadherin, N-cadherin, Snail, and Vimentin) was measured using Western blot and qPCR. The result showed that, compared to EVC, overexpression of CD24^WT^ reduces the expression of E-cadherin and increases the expression of N-cadherin in both HCT116 and SW480 **(Figure 4A, D)**. This phenomenon is well-described in the literature as the “cadherin switch” and is associated with EMT [24]. Conversely, transfection with CD24^N[36,52]Q^ showed a complete failure to induce the cadherin switch in SW480 and a markedly diminished effect in HCT116. Complementing the cadherin switch, Snail and Vimentin levels were increased in HCT116 and SW480 by forced expression of CD24^WT^. The level of these markers was lower in the forced expression of CD24^N[36,52]Q^ vs CD24^WT^.

**Figure 4.**
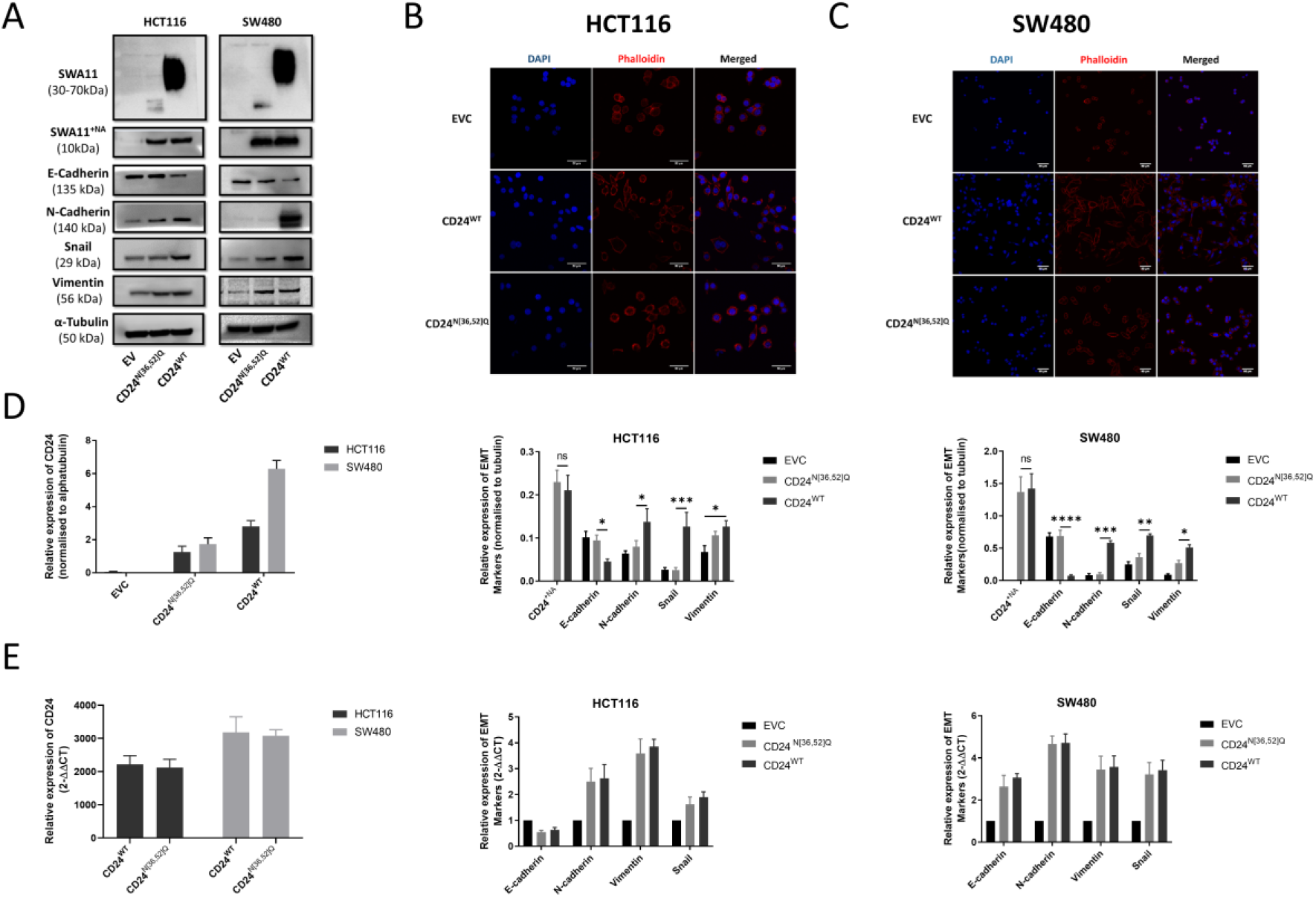
CD24 N-glycosylation is associated with EMT phenotype alteration and cadherin switch in CRC cells. Expression of CD24 along with EMT-related biomarkers (E-cadherin, N-cadherin, Snail, and Vimentin) was determined by WB post-transfection of HCT116 and SW480 cells with CD24 ^WT^ and CD24^N[36,52]Q^. The cropped blots are used in the figure. The membranes were cut before to exposure so that only the portion of gel containing the desired bands would be visualized. The samples derive from the same experiment and the blots were processed in parallel (**A**). Cytoskeleton F-actin staining at ×20 magnification shows CD24^WT^ involvement in EMT morphological changes in HCT116 (**B**) and SW480 (**C**) and this restrained in CD24^N[36,52]Q^. The quantitative densitometry analysis of western blot for CD24 and EMT proteins normalised to alpha-tubulin **(D)**. RT-qPCR analysis of CD24 and EMT biomarkers in CD24^N[36,52]Q^ showed no change in the mRNA levels compared to CD24^WT^ overexpressed cells **(E)**.

We have unpublished data suggesting that CD24 may alter the target molecules’ levels at both the transcriptional and post-transcriptional level. The mRNA levels of these targets were, therefore, evaluated by quantitative PCR. In both cell lines, the mRNA levels of N-cadherin, Snail and Vimentin were increased by transfection with both CD24^N[36,52]Q^ and CD24^WT^ when compared to EVC. There was, however, no difference in levels of mRNA expression seen between the two constructs. Similarly, no difference was seen in the levels of E-cadherin mRNA following transfection of these constructs but, in SW480, the levels of E-cadherin mRNA were higher than that in HCT116 **(Figure 4E)**.

EMT is a morphological change whereby tumour cells lose epithelial properties to gain a mesenchymal appearance. To investigate EMT further, phalloidin staining for filamentous-actin (F-actin) was performed. In comparison to cells transfected with EVC, cells transfected with CD24^WT^ acquired spindle-like features, and phalloidin staining showed actin polymerization and the formation of actin filaments in the cytoplasm. In contrast, cells overexpressing CD24^N[36,52]Q^ maintained a rounded appearance, with only a few cells acquiring an elongated shape. However, the staining with phalloidin was similar in CD24^N[36,52]Q^ vs CD24^WT^ suggesting that, despite the loss of the N-glycosylation sites, CD24 was still able to induce actin polymerisation **(Figure 4B, C)**. Further studies are warranted to investigate actin polymerisation through the quantification of globular (G)- and F-actin cellular fractions through Triton-X.

## Discussion

CD24 is a putative gene expressed in various cancer types, including CRC, where its overexpression is defined as a key early event in the development of CRC [25]. Glycosylation in many proteins plays a significant role in normal biological processes and cancer development, including tumour cell proliferation, invasion, angiogenesis, and metastasis [26]. Changes in a specific *N*-linked glycoprotein are associated with the development of many diseases; and thus, it is crucial to study the *N*-linked glycoproteins to facilitate understanding of disease and help develop effective treatments [27,28].

The structure of CD24 shows two putative N-glycosylation sites located around codon 36 and codon 52. In this study, we investigated the role of these sites in mediating the known functions of CD24, such as cell motility and EMT. Firstly, we proved that these sites were bona-fide N-glycosylation sites. At both sites, we substituted the asparagine with the glutamine **(Figure S2)**, thereby removing the N-glycosylation target motif. This substitution would not affect the other *O*-glycosylation sites and would not create any new *O*-glycosylation sites as glutamine cannot bind to *O*-glycans [29]. Two constructs were created in which each N-glycosylation site was removed individually (CD24^N36Q^ and CD24^N52Q^), and a third construct in which both sites were removed (CD24^N[36,52]Q^). Western blotting confirmed that each of the mutant constructs was expressing the protein, although the banding patterns were different from the protein expressed by the wild-type construct (CD24^WT^). Plasmid transfection efficiency and expression of core protein were similar between the different constructs leading us to conclude that the variability in the Western blot banding patterns was due to variation in the glycosylation. The wild-type protein showed the classical smeared band. Protein from each single-mutation construct (CD24^N36Q^ and CD24^N52Q^) showed less smearing and quicker running speed than the wild-type protein; these features are consistent with reduced glycosylation. The single-mutation proteins bands were very similar, suggesting they had similar levels of glycosylation. Protein from the double mutant construct (CD24^N[36,52]Q^) contains only O-glycosylation sites, and the smearing of the bands suggested that there was glycosylation. However, even though there are 16 putative glycosylation sites, the banding pattern suggests that the overwhelming majority of glycosylation of CD24 lies at the N-glycosylation sites. Our observations are the first description of this pattern of glycosylation in CRC.

Having shown that the putative N-glycosylation were indeed true glycosylation sites, we next investigated whether N-glycosylation was relevant to the known cellular function of CD24, e.g., regulation of EMT, cell migration, and cell invasion in CRC cells, as these are crucial steps for tumour metastasis [30]. All assays (transwell migration and wound healing for cell migration and transwe**l** migration through Matrigel for invasion, immunofluorescence with phalloidin staining for EMT) showed similar results, i.e., the activity of the single mutants was significantly less than that of the CD24^WT,^ thereby suggesting each of the sites contributed to these functions. Furthermore, the effect seemed to be additive, as the activity of the double mutant (CD24^N[36,52]Q^) was significantly less than the effect of a single mutation (leading to the near-total abrogation of the ability to CD24 to induce some of these processes) when both N-glycosylation sites were mutated. To our knowledge, this is the first report of the functional importance of the N-glycosylation in CD24. Our data were complemented by Western blots showing that loss of N-glycosylation sites resulted in the failure of the cadherin switch and a reduced level of induction of the motility-related biomarkers Snail and Vimentin. The mRNA’s evaluation showed similar levels of induction of gene transcription of N - cadherin, Snail, and Vimentin by both the wild-type CD24 and double mutant CD24^N[36,52]Q^. This would suggest that the changes observed in protein expression were mostly due to post-transcriptional modifications.

Our results are comparable to other studies that identify the potential role of N-glycosylation in inducing migratory and invasion capabilities of tumour cells [31-33]. The exact mechanism through which the N-glycosylation sites induce these changes is uncertain, but we would conjecture that the glycosylated asparagine residues at codons 36 and 52 are able to form complexes with and activate target cell surface molecules. The additive effects could be explained by each glycosylated residue contributing a certain amount to the total avidity for the target molecules. Since both transcriptional and post-transcriptional effects are seen, it is likely that several different pathways are activated. This would fit with the multiple binding partners described for CD24 [6].

EMT is associated with the acquisition of cell motility [34]. We and others previously have demonstrated CD24 involvement in EMT of CRC cells [6,35]. In line with the cell motility studies, immunofluorescence studies showed that cells transfected with CD24^WT^ acquired a spindle-shaped mesenchymal phenotype (compared to cells transfected with EVC), and this was accompanied by actin polymerization. In cells transfected with the double mutant, CD24^N[36,52]Q^, the cells retained a rounded epithelioid phenotype similar to that seen in cells transfected with EVC. The only inconsistency in our data was that cells transfected with CD24^N[36,52]Q^ also showed actin polymerization at a level similar to that seen with CD24^WT^. Thus, it is apparent that CD24 can induce actin polymerization and be mediated by mechanisms independent of N-glycosylation. It is uncertain what these mechanisms, whether they are related to the residual low-level O-glycosylation and how they are coordinated with the N-glycosylation-dependent mechanisms of motility induction.

Previously we have shown that CD24 has no influence on proliferation in CRC cell lines [5]. Therefore, it was surprising to observe that mutation of either of the *N*-glycosylation sites resulted in increased proliferation in both HCT116 and SW480. As with the study, the size of the effect was similar for each mutation but, when combined together in the construct CD24^N[36,52]Q^, the effects appeared additive. There is obviously a gain-of-function acquired through the removal of the N-glycosylation sites, which suggests that some signalling pathways are activated which are not activated by wild-type CD24. We would conjecture that loss of N-glycosylation may allow the CD24 to form complexes with and activate cell surface molecules (such as growth factor receptors) from which it would ordinarily be shielded by the sugar residues around codon 36 and 52. If it is a true biological phenomenon, it may be related to the role of CD24 as an inducer of stemness [19,23]. Stem cells often have proliferative quiescence, and it is possible CD24 induces this through N-glycosylation. Alternatively, it could be an artefact of our experimental system. Either way, our data warrant further investigation to identify the targets which are leading to this observed stimulation of proliferation and identifying whether CD24 colocalizes with any growth factor receptor.

To summarize, we have confirmed, in two different CRC cell lines, that the putative *N*-glycosylation sites of CD24 around codon 36 and codon 52 are true targets for glycosylation. For the first time, these sites have been shown to be functionally active mediators of CD24-induced EMT and cell motility (both cell migration and cell invasion) in CRC. The function of the N-glycosylation sites is mediated through both transcriptional and post-transcriptional mechanisms, although actin polymerization may be mediated through N-glycosylation independent pathways of CD24. Unexpectedly, the N-glycosylation sites may also play a role in cell proliferation. These results suggest that CD24 *N*-glycosylation may play a role in progression and metastasis in CRC. Further studies are necessary to elucidate the mechanisms of CD24 activity in CRC and explore whether it could be a target for therapy.

## Supplementary Materials

All supplementary materials have deposited in a separate file.

## Authors’ contributions

Z.H performed all the experimental work, the data analysis and wrote the manuscript. T.R assisted with cell culture and the analysis of the data. A.A assisted with the design of the primer sequences. W.D assisted with the design of the primer sequences and revised the manuscript. H.O assisted in result interpretation. C.H and K.G edited and reviewed the manuscript. M.I designed the study and edited and revised the manuscript. All authors have read and agreed to the published version of the manuscript.

## Institutional Review Board Statement

Not applicable

## Informed Consent Statement

Not applicable

## Ethics approval and consent to participate

Not applicable

## Data Availability Statement

The data presented in this study are available from the corresponding author on reasonable request.

## Acknowledgements

The author would like to express the deepest appreciation to all colleagues in the Pathology Research Group and the Division of Cancer and Stem cells for their kind support and assistance.

## Supplementary Material

### Supplementary tables

**Table S1:**
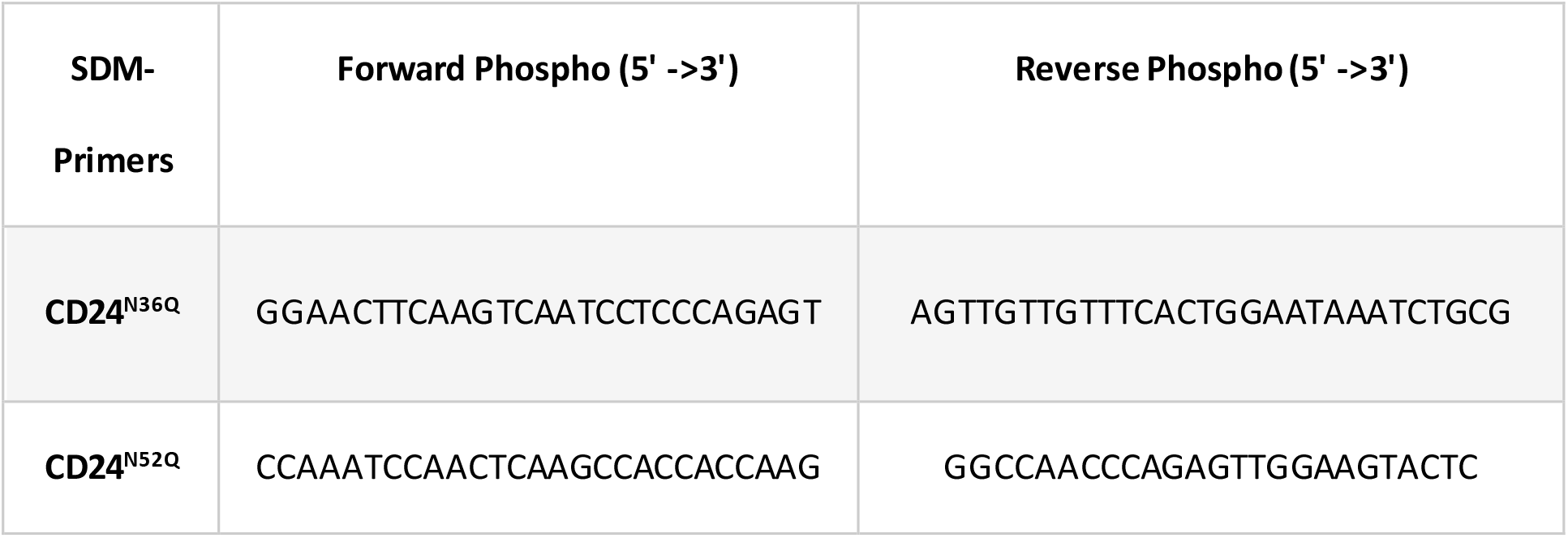
Primer sequences used for site-directed mutagenesis (SDM)

**Table S2:** Details of relevant primary and secondary antibodies used for Western blot

| Antibody | Provider | Species | Diluent | Ab<sub>1</sub> dilution | Ab<sub>2</sub> dilution |
| --- | --- | --- | --- | --- | --- |
| **CD24**<br>**(SWA11)** | Prof. Altevogt P<br>(DKFZ, Heidelberg) | Mouse | 5% Skimmed<br>Milk+0.1%<br>Tween PBS | 1:1000 | 1:5000 |
| **E-cadherin** | Cell signalling | Rabbit | 5% BSA+0.1%<br>Tween TBS | 1:1000 | 1:5000 |
| **N-cadherin** | Cell signalling | Rabbit | 5% BSA+0.1%<br>Tween TBS | 1:1000 | 1:5000 |
| **Snail** | Cell signalling | Rabbit | 5% BSA+0.1%<br>Tween TBS | 1:000 | 1:5000 |
| **Vimentin** | Cell signalling | Rabbit | 5% BSA+0.1%<br>Tween TBS | 1:000 | 1:5000 |
| **Tubulin** | Abcam | Mouse | 5% Skimmed<br>Milk+0.1%<br>Tween PBS | 1:000 | 1:5000 |

**Table S3:**
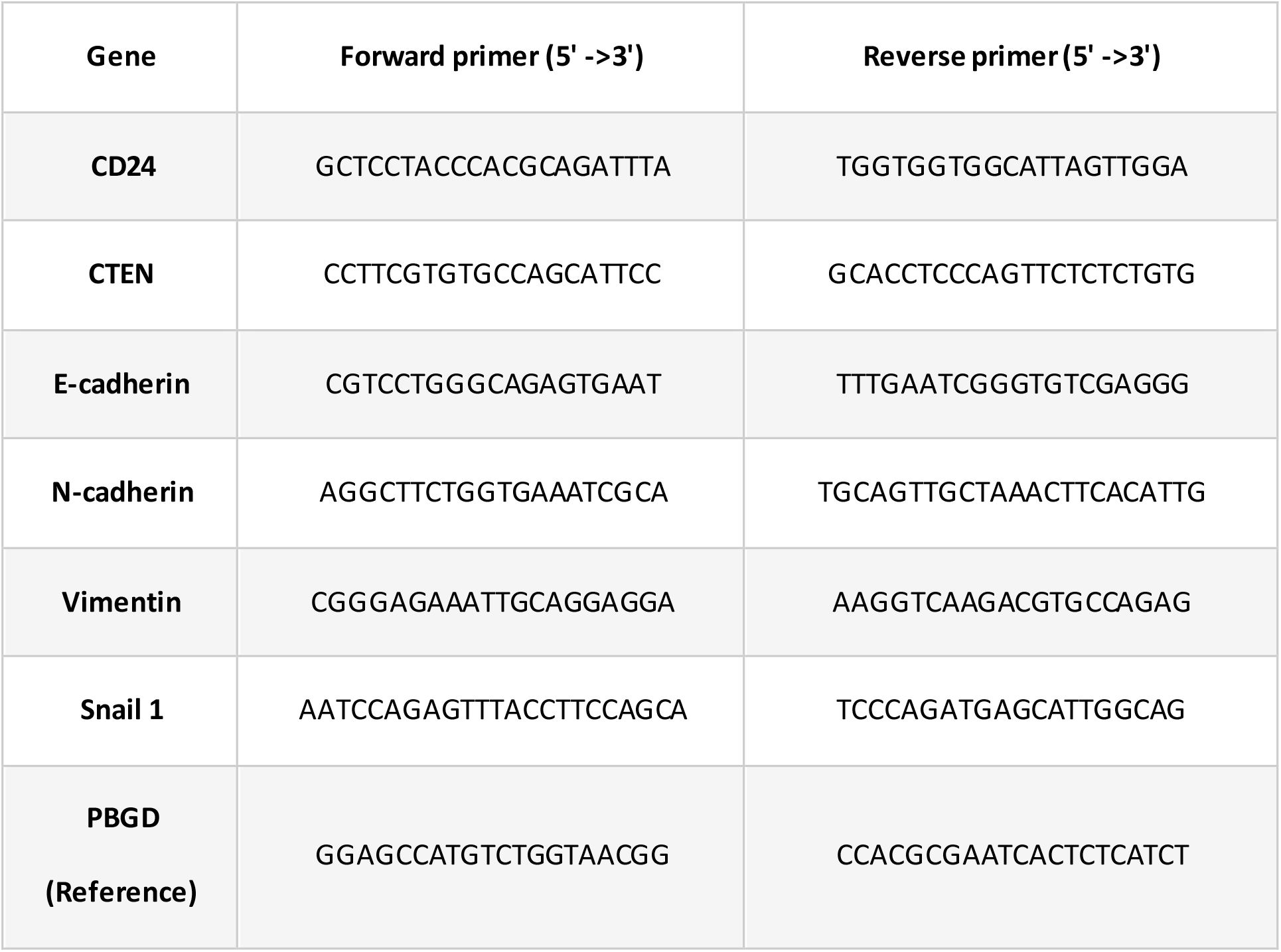
Primer sequences used for quantitative real-time PCR

### Supplementary Figures

**Figure S1:**
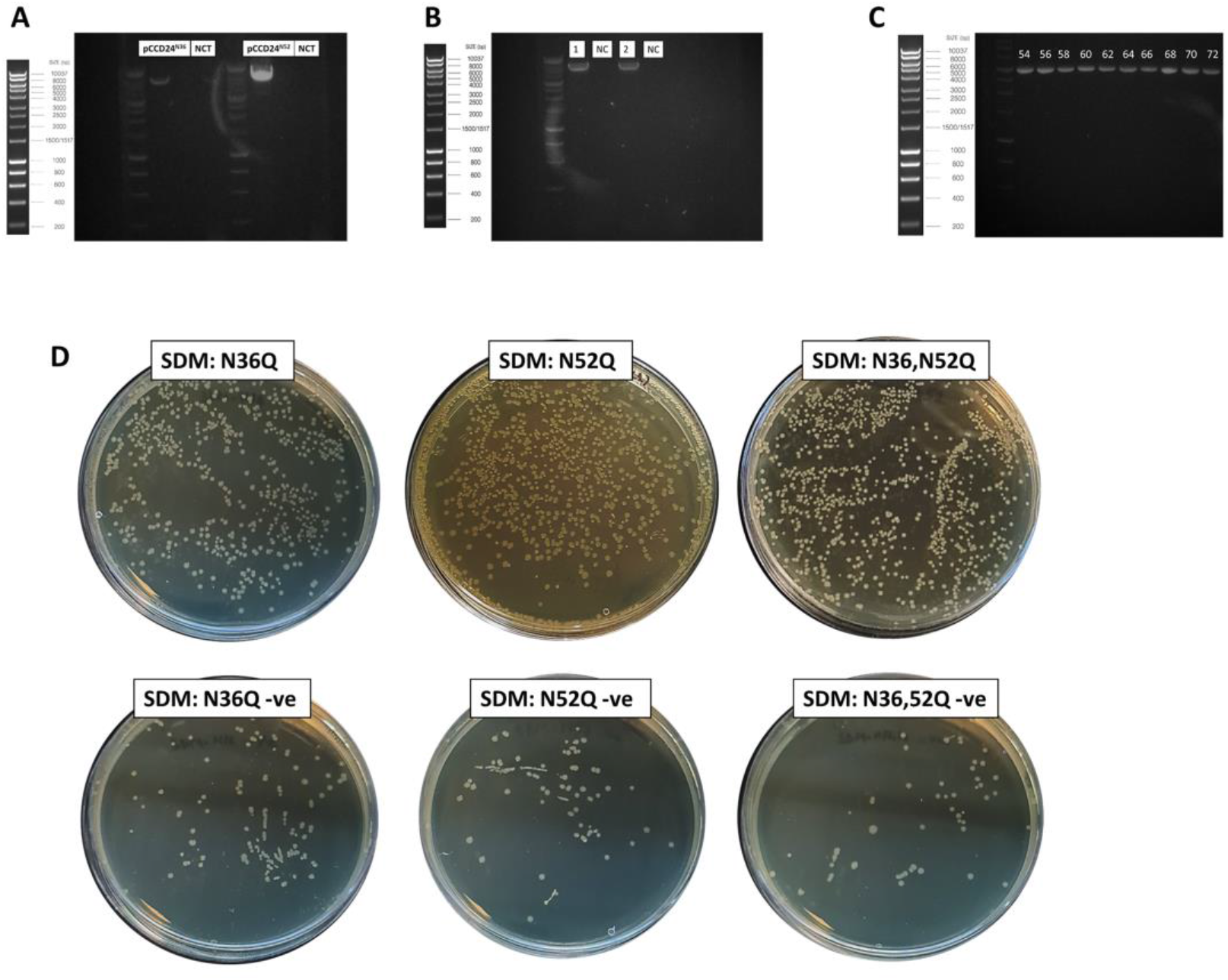
Determination of SDM confirmation in CD24^N36Q^, CD24^N52Q^, and CD24^N[36,52]Q^ Plasmids with Gel Electrophoresis and Colony Formation. Gel electrophoresis of the SDM products produced bright bands at the correct molecular size. **(A, B)** Primers were amplified at 68°C. No band was observed in the no-template control lane. **(C)** Site-directed mutagenesis was performed on two sets of back to back primers—a gradient PCR for optimising the annealing temperature designed primers. Showing the correct molecular weight of 5747kb. **(D)** Bacterial colony formation in mutant clones (CD24^N36Q^, CD24^N52Q^, and CD24^N[36,52]Q^) compared to the negative control, showing successful ligation. All experiments performed in triplicate.

**Figure S2:**
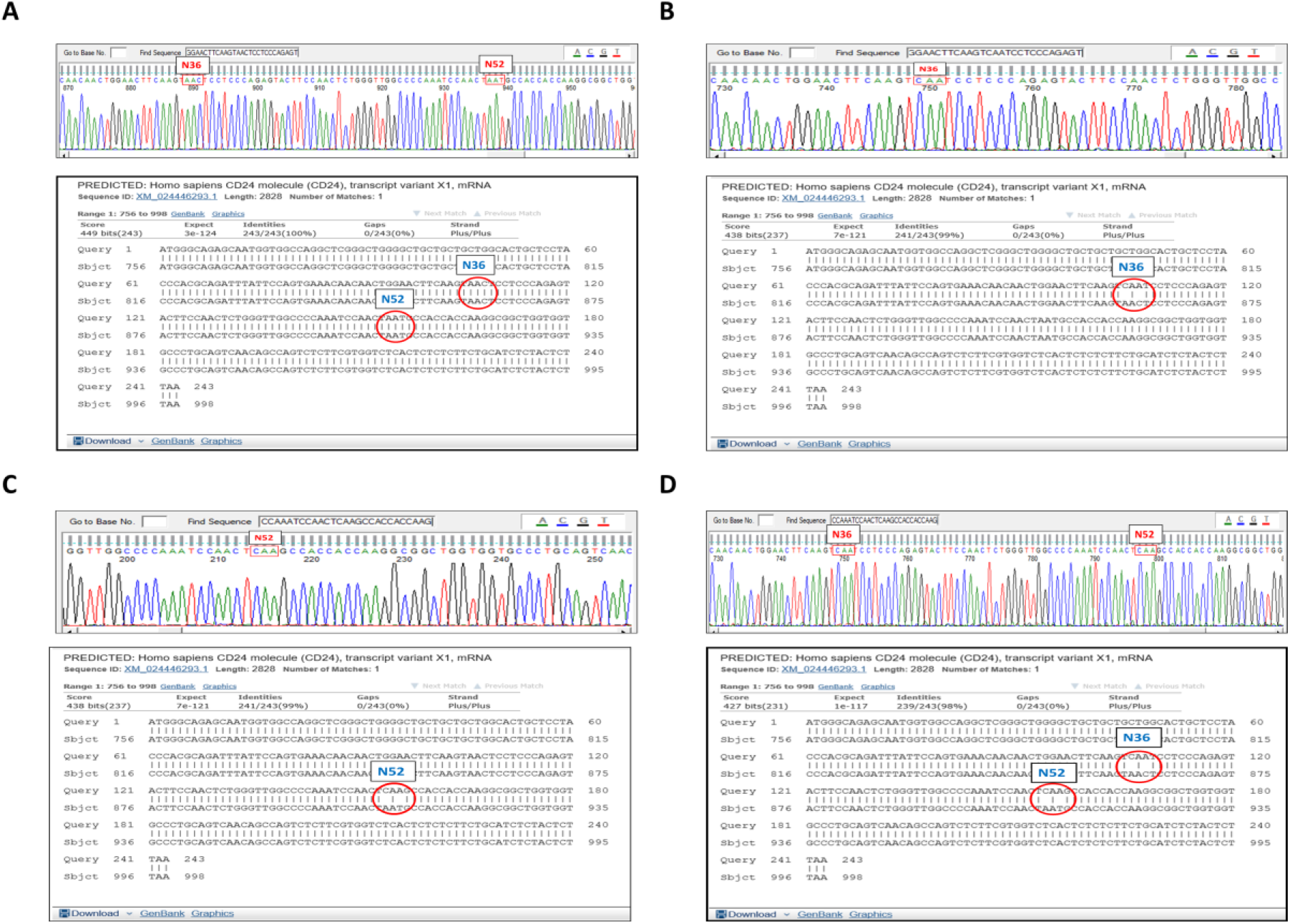
Gene sequencing analysis using Finch software and alignment tool (Bast2). The presence of the CD24 gene without any mutation at both indicated codons N36 and N52 with 100% of nucleotides homology (highlighted) **(A)**. The single-mutant clones showed the desired mutation at N36, i.e. AAC to CAA **(B)**, and N52, i.e. AAT to CAA **(C)**, with 99% homology in nucleotides sequence (top). The double-mutant clone showing the mutation at both N36 and 52 simultaneously, and the alignment tool showed 98% similarity in nucleotides sequence) **(D)**. All sequencing databases for CD24^N36Q^, CD24^N52Q^, and ^CD24N[36,52]Q^ have been deposited to the GeneBank and will be released as soon as possible. The GenBank accession numbers for mutant constructs are MW694788, MW694789 and MW694790, respectively.

**Figure S3:**
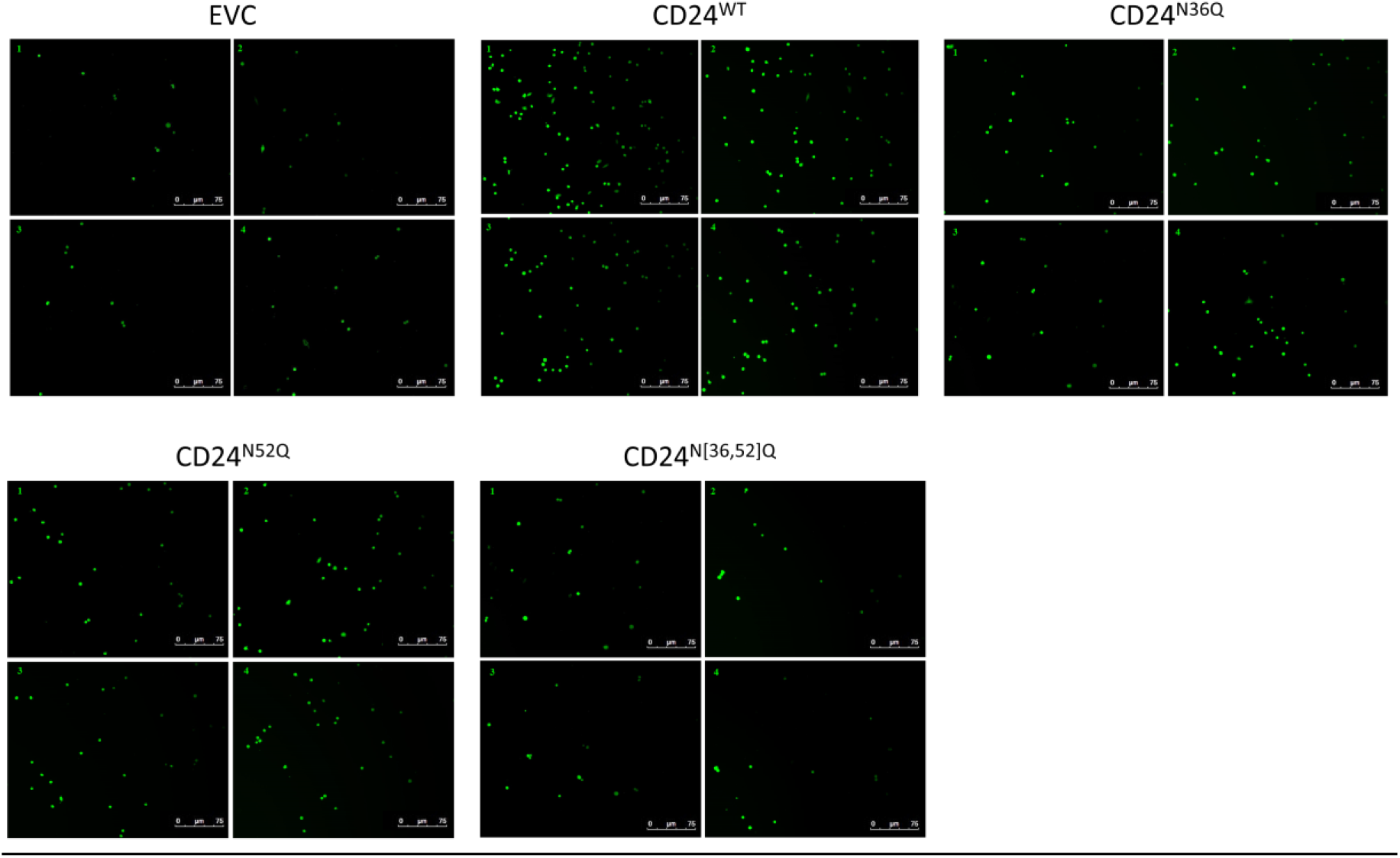
Representation of the transwell assay analysis of migrating and invasive cells. Four random microscopic fields per well were taken at ×10 magnification for EVC, CD24, and CD24 mutant constructs (CD24^N36Q^, CD24^N52Q^, and CD24^N[36,52]Q^). The number of cells was counted manually using ImageJ (National Institutes of Health, Bethesda, MD) and analysed by taking the average of the four fields per wach well. Three exprimental replicates was performed. Abbreviation: EVC: Empty vector control.

## Supplementary methods

### Method S1: Flow cytometry

Transfection efficiency for cells overexpressed with CD24^WT^ and mutant clones was determined using FACS analysis. Twenty four hours post-transfection, the cells were detached using Trypsin-EDTA and resuspended in sterile-filtered PBA (Phosphate buffered saline +1% bovine serum albumin). The cell suspension washed twice using PBA buffer by centrifugation for 5min at 300g. The cell was counted using an automated cell counter, and the volume of cells in PBA was adjusted at 2×10^6^/ml and pelleted by centrifugation at 300g for 5 min. Then the cell pellet resuspended in 100μl of PBA and incubated with CD24 (SWA11) antibody (1:100) for 60 min on ice. The antibody was washed twice by adding 1ml of PBA and centrifuged for 5 min at 300g. The cell pellet resuspended in 100μl of PBA and Alexa Fluor 488 goat anti-mouse conjugated secondary antibody (Invitrogen) was added. The cells were incubated for 30min on ice in the dark and then washed twice with PBA. The supernatant was removed, the pellet was vortexed and resuspended with 500μl of 4% formaldehyde. The transfection efficiency was assessed by determining the ratio of cells overexpressing CD24 using a Beckman Coulter FC500 Flow Cytometer (Beckman Coulter) located in the Flow Cytometry Facility at Medical School, Queen Medical Centre. The data were analysed using Kaluza analysis 2.1 software (Beckman coulter Inc.), in which the cells were gated for identification of the CD24+ population using non-transfected cells as a control.

### Method S2: N-glycosylation digestion

Transfection of the modified construct of CD24, which was less glycosylated than the wild type CD24, resulted in variation in the molecular weight on western blot. Therefore, the glycosylation of CD24 was removed using an N-glycosylation digestion mix that contains a combination of α2-3, 6, 8 Neuraminidase and Peptide -N-Glycosidase F (PNGase F) (New England BioLabs, UK). According to the provided instructions, the reaction was prepared in a reaction volume of 20 μl as follows: 25 μg of the protein was added to 1 μl of glycoprotein denaturing buffer (10X) H2O to make 10 μl reaction volume. Next, the protein was denatured by heating the reaction at 100°C for 10 minutes, followed by ice - chilled and centrifugation for 10 seconds. After that, the total reaction volume was made to 20 μl by adding 2 μl GlycoBuffer (10X), 2 μl 10% NP-40 and 6 μl of water. Following this, 1μl PNGase F and 1μl α2-3,6,8,9 Neuraminidase (2μl/1μg) were mixed with the 20 μl-volume reactions and incubated at 37°C overnight. After incubation, the enzyme was deactivated by incubating the protein at 68°C for 10 minutes. The mixture then was run on an SDS-PAGE gel.

### Method S3: Complementary DNA (cDNA) synthesis

The cDNA was synthesised from RNA templates as follows: 15μL reverse transcription master mix per reaction was prepared by adding 0.5μg random hexamers (Thermo Fisher Scientific) to 1-2µg RNA, and nuclease-free water, incubated for 5mins at 70°C, then by 5mins at 4°C. Followed by making 10μL/ reaction of the bulk mixture containing: 5μL M-MLV-RT buffer (5X Promega), 1.25μL dNTPs (10mM, Promega), 1µL M-MLV-RT enzyme (Promega), 25U RNase inhibitor (0.625μL) (Promega), made up to 10μL with distilled water. 10μL of the bulk mixture was added to the 15μl M-MLV template: hexamer mix to a total volume of 25μl and incubated for 1hr at 37°C, then 10mins at 95°C.

